# Duplication of the anteroposterior body axis in postembryos of the *Eratigena atrica* spider

**DOI:** 10.1101/2024.02.14.580276

**Authors:** Teresa Napiórkowska, Julita Templin, Paweł Napiórkowski

**Affiliations:** Department of Invertebrate Zoology and Parasitology, Faculty of Biological and Veterinary Sciences, Nicolaus Copernicus University in Toruń, Toruń, Poland; Department of Hydrobiology, Faculty of Biological Sciences, Kazimierz Wielki University in Bydgoszcz, Bydgoszcz, Poland

**Keywords:** AP axis duplication, Alternating temperatures, Body anomalies, Cumulus, Spider, Teratology

## Abstract

Many factors, including temperature, can affect the embryonic development of spiders, leading to a variety of deformities in their body structure. Early studies on teratological changes in spiders were aimed mainly at describing the morphology of affected individuals. Now, with new tools and molecular techniques used to carry out a functional analysis of genes involved in body segmentation and leg formation, the explanation of these defects is finally possible. In our new experiment on spiders collected in 2022/2023 breeding season, we obtained 87 postembryos with various body defects by applying alternating temperatures (14^0^C and 32^0^C) for the first 10 days of embryonic development. The most common anomaly, i.e. oligomely, occurred in 70 individuals, while other defects (schistomely, symely, polymely, heterosymely) were noted sporadically. Partial prosoma duplication affected only six embryos. In the context of recent advances in molecular spider embryology we conclude that recorded anomalies may be connected with the suppression or erroneous expression of certain genes of body segmentation and appendage formation, and by disturbances in the formation and migration of the cumulus, which leads to an axis duplication phenotype.

**Summary statement:** This article presents a rare anomaly - bicephaly, which appeared in spider postembryos as a result of thermal shocks applied during embryogenesis.

## INTRODUCTION

Many factors, including temperature, can significantly affect the embryonic development of arthropods (e.g. Juberthie, 1962, 1968; Itow, 1982, Buczek, 2000), leading to a range of body defects of varying severity and complexity. Literature abounds in reports of deformities of all body segments, appendages and parts of the external skeleton in crustaceans, insects, myriapods, and chelicerates (Itow and Sekiguchi, 1980; Shelton et al., 1981; Ćurčić et al., 1983; Spanò et al., 2003; Köhler et al., 2005; Mattoni, 2005, Kamaruzzaman et al., 2006; Mąkol and Łaydanowicz, 2006, Asiain and Márquez, 2009; Eeva and Penttinen, 2009; Gregati and Negreiros-Fransozo, 2009; Fernandez et al., 2011; Kozel and Novak, 2013; Scholtz and Brenneis, 2016). Much more complicated and relatively less frequent are duplications of certain body parts or even the entire body, referred to as conjoined twins. These are characterized by a partial or complete duplication of the anteroposterior body axis and show various degrees of fusion. Scientists traditionally distinguish between conjoined twins with double anterior structures such as the head, referred to as *Duplicitas anterior* (*DA*) or twins with double posterior regions, referred to as *Duplicitas posterior* (*DP*). The double embryos each of which has its own complete anteroposterior body axis are called *Duplicitas completa* (*DC*) (Scholtz, 2020). These anomalies were investigated, e.g. by Scholtz et al. (2014) in *Amarinus lacustris*, Rudolph and Martinez (2008) in *Virilastacus rucapihuelensis*, Harzsch et al. (2000) in *Homarus americanus* and Jara and Palacios (2001) in *Aegla abtao*. Duplication of body parts in chelicerates have also been studied. Estrada-Peňa (2001) described three specimens of *Rhipicephalus sanguineus* with different degrees of twinning *DP*-type, Itow and Sekiguchi (1979) obtained many embryos of *Tachypelus tridentatus* with *DA*, *DP* and *DC* malformations, Matthiesen (1979) presented an embryo of the *Tityus cambridgei* scorpion with an anterior body duplication (*DA* malformation), and Seiter and Teruel (2014) described two cases of metasoma duplication (*DP*) in buthid scorpions. In the literature there are also a few examples of anterior body duplication in spiders. Rempel (1954) described two *Latrodectus mactans* embryos with prosomal duplication, *Tegenaria atrica* with two heads were also described by Mikulska and Jacuński (1970, 1971), who obtained this type of anomaly by increasing temperature during embryo incubation. The morphology of the prosoma of bicephalic *Tegenaria atrica* has been investigated on several occassions by Jacuński and Templin (2003), Templin et al. (2009), Napiórkowska and Templin (2017a) and Napiórkowska et al. (2021). All bicephalic postembryos were obtained in experiments using alternating temperatures, which not only led to the duplication of the prosomal front but also, occasionally, caused a number of additional changes, complicating the morphology of this body part to a large extent. In several cases, the structure of the central nervous system was also examined (Jacuński and Templin, 1992; Jacuński et al., 2002; Napiórkowska et al., 2010a, 2016c). Since *DA* anomaly is characterized by considerable diversity (includes cases of two fully developed heads, one fully-developed head and one incomplete head with additional complications resulting from the fusion of the chelicerae and/or pedipalps), the structure of the central nervous system can be affected. First attempts were also made to explain the causes of duplication of the body front in spiders (Jacuński and Templin, 2003; Templin et al., 2009).

Duplication of the body front in arthropods, including spiders, seems to be rare. The results of many teratological studies using alternating temperatures indicate that it affects only a few spiders in one reproductive season (Jacuński et al., 2004; Napiórkowska and Templin, 2017b, Napiórkowska et al., 2016a,c, 2021) thus constituting only a small percentage of all developmental anomalies in postembryos. On the other hand, oligomely (the absence of one or more appendages) is the most common (Jacuński et al., 2005; Napiórkowska and Templin, 2013; Napiórkowska et al., 2013, 2015, 2016b). Other anomalies, such as symely (fusion of contralateral appendages), schistomely (bifurcation of appendages), heterosymely (fusion of ipsilateral appendages), polymely (presence of one or more additional appendages) and so-called complex anomalies (two or more anomalies occurring simultaneously) are recorded sporadically (e.g. Napiórkowska et al., 2010b, 2017; Napiórkowska and Templin, 2018). Many of them, including appendages on a pedicel, described by Jacuński (1971, 1984), Jacuński and Templin (1991) and Napiórkowska et al. (2023), are of great interest for evo-devo research.

Early teratological studies of spiders focused mainly on the morphology of affected individuals. At present, owing to the availability of new tools and techniques, both the analysis of deformities and the explanation of their causes are less challenging. As a consequence, a functional analysis of genes involved in body segmentation and leg formation in arthropods is possible. The development of RNA interference (RNAi) technique, in situ hybridization and immunolabeling resulted in the dynamic evolution of research on the expression of many developmental regulatory genes during spider embryogenesis (Stollewerk et al., 2003; Schoppmeier and Damen, 2005; Oda et al., 2007; McGregor et al., 2008; Schwager et al., 2009; Pechmann et al., 2011; Khadjeh et al., 2012; Oda and Akiyama-Oda, 2019; Heingård and Janssen, 2020; Setton and Sharma, 2021). Based on this, we can safely assume that at least some of the observed anomalies in spiders are caused by the erroneous suppression or expression of segmentation or appendage patterning genes (e.g. Khadjeh et al., 2012) or by disturbances in early embryogenesis, e.g. in the formation and migration of the cumulus (the organizing centre of early spider embryos), which moves from the centre of the germ-disc towards the periphery of the disc and activates the BMP signalling pathway in ectodermal cells that are close to the cumulus (e.g. Akiyama-Oda and Oda 2003, 2006; Hilbrant et al., 2012; Oda and Akiyama-Oda, 2008; Pechmann et al., 2017). This activation of the BMP signalling pathway in a subset of germ-disc cells is essential to break the radial symmetry of the disc and to initiate the formation of the dorsal field and the bilateral symmetry of the spider embryo (McGregor et al., 2008; Oda et al., 2020; Schwager et al., 2015). Any disturbances at this stage of embryo development may lead to an axis duplication phenotype (Pechmann, 2020).

Through many years of research, we have obtained a relatively large number of postembryos with body defects. Our study was aimed at exploring the diversity of developmental anomalies induced by the variable temperature protocol, with particular attention to partial prosoma duplication observed only in six *E. atrica* postembryos. Furthermore, we attempted to interpret these anomalies in terms of disturbances in cumulus formation and migration, focusing on functional studies on spiders and other invertebrates.

## RESULTS

In the 2022/2023 breeding season we obtained approximately 9,400 embryos, half of which constituted the control group. In postembryos that left eggshells, no anomalies were found on the prosoma or the opisthosoma. All individuals had six pairs of properly segmented appendages (Fig. 1) and a properly developed spinning apparatus. Embryo mortality in this group was 9%.

**Fig. 1.**
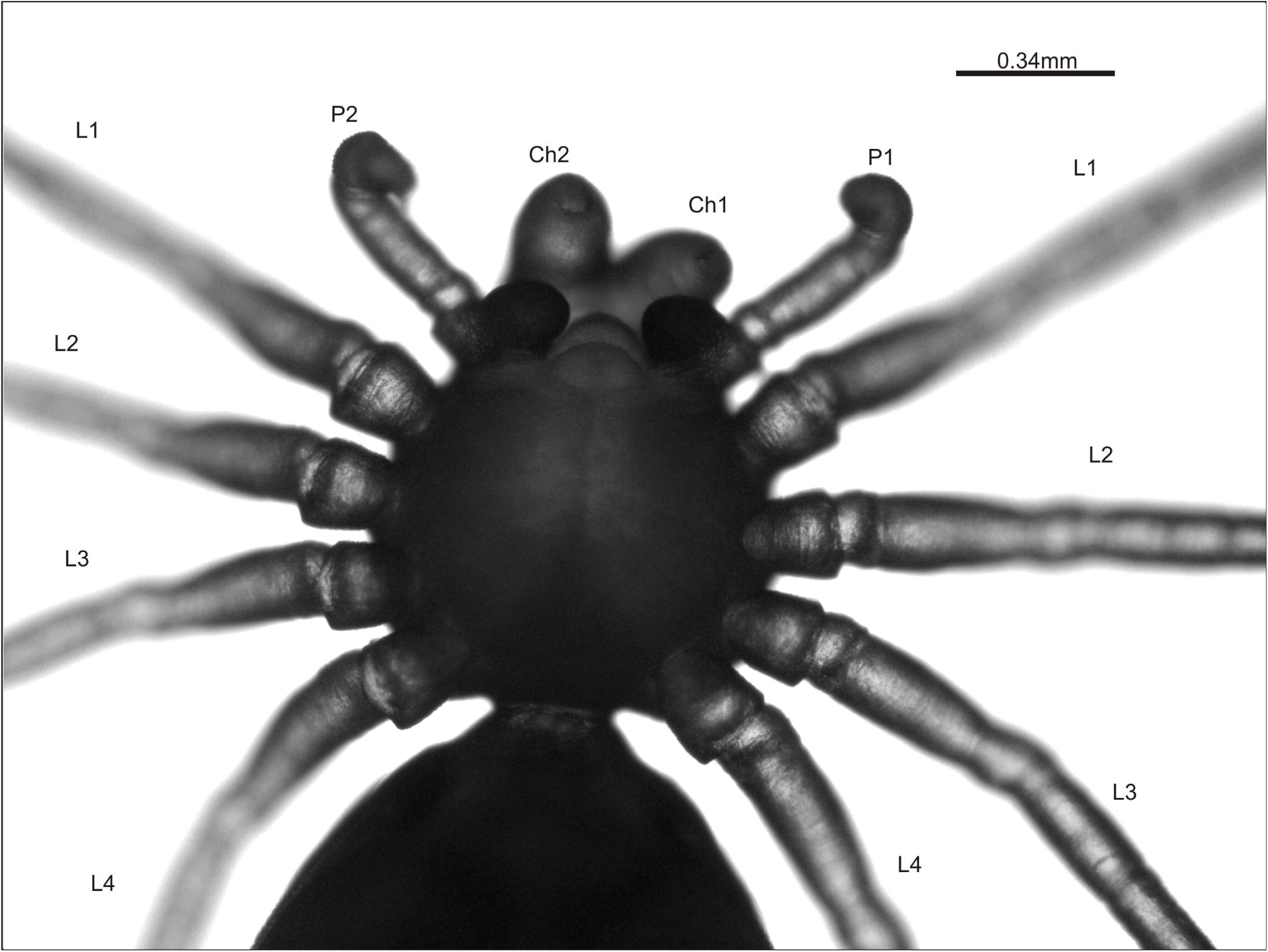
Normally developed *Eratigena atrica* postembryo from the control group (ventral view). Ch, chelicera; L1-L4, walking legs 1-4; P, pedipalp.

In the experimental group embryo mortality was much higher, amounting to 38%. Out of approximately 2,900 postembryos that left eggshells on their own, 87 (about 3%) had body defects (Table 1), mainly on the prosomal appendages. The vast majority of the affected postembryos lacked an appendage/appendages (oligomely), and in almost 57% of the total number of teratogenically altered individuals it was a walking leg/walking legs that was/were missing. On the other hand, schistomely, symely, polymely, and heterosymely were only recorded in several individuals. The “Other abnormalities” group includes spiders with considerably shorter appendages on the prosoma and protuberances of different sizes and shapes. Only two individuals had changes in the spinning apparatus (oligomely) on the opisthosoma. However, the most interesting group includes postembryos with significant changes in the front of the prosoma. All these individuals, classified as individuals with a duplicated front part of the prosoma, are shown in Fig. 2A-F.

**Fig. 2.**
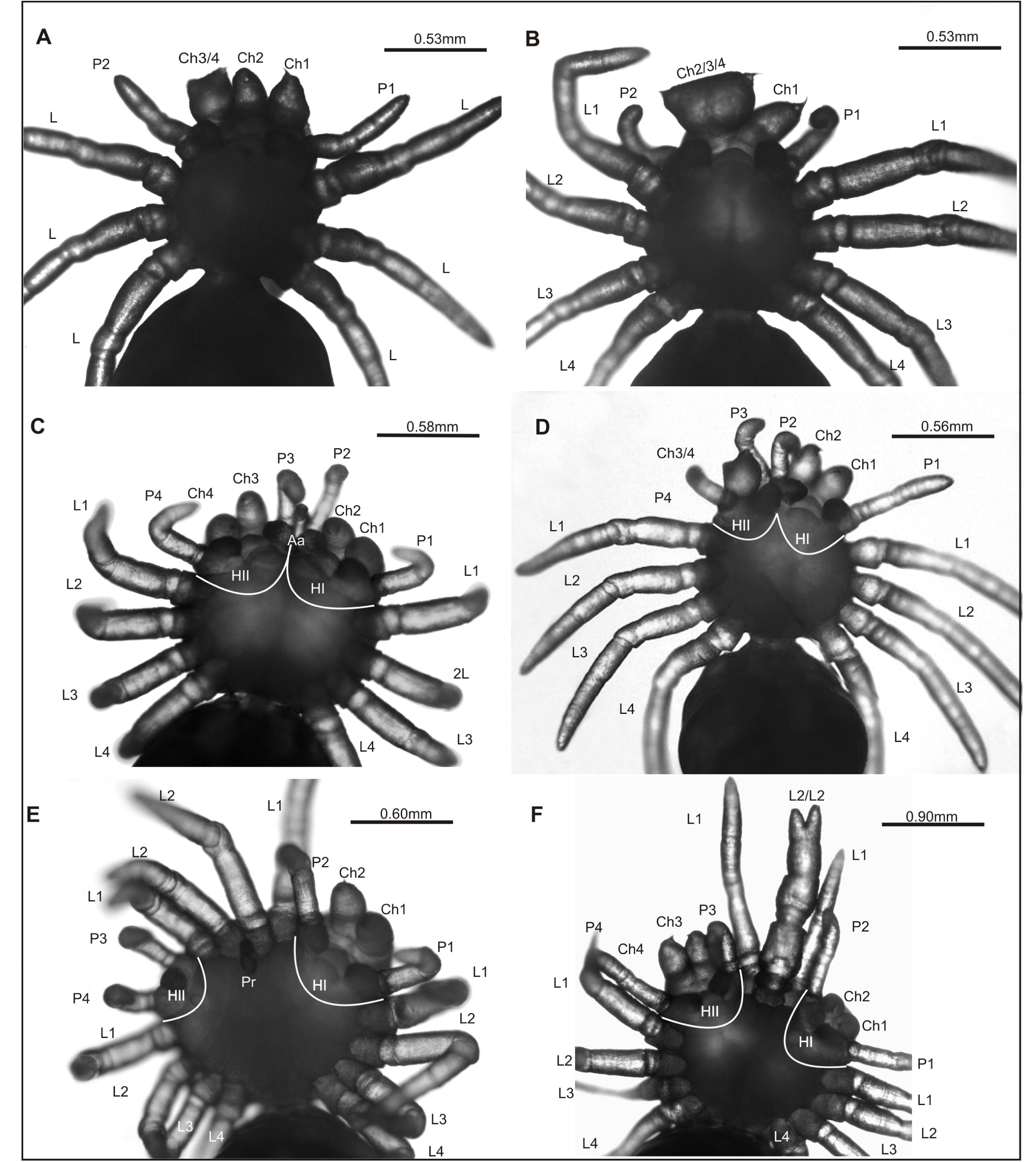
*Eratigena atrica* postembryos with anteroposterior (AP) body axis duplication (ventral view). A: postembryo with two normally developed chelicerae (Ch 1, Ch2) and additional partially fused chelicerae (Ch 3/4), lacking two walking legs, one on the right and one on the left side of the prosoma; B: postembryo with a well-developed chelicera (Ch1) and three chelicerae partially fused (Ch 2/3/4); C: bicephalic postembryo with a normally developed heads (HI and HII) and with an additional, shortened appendage (Aa); D: bicephalic postembryo with a normally developed left head (H I) and a right head (H II) with partially fused chelicerae (Ch 3/4); E: bicephalic postembryo with a normally developed left head (H I) and right head without chelicerae (H II) but with four walking legs (L1 and L2) between the heads; F: bicephalic postembryo with two normally developed heads (H1 and H II) and with four walking legs between the heads (L1 and L2), two of which are fused (L2/2); white lines indicate the heads of bicephalous postembryos.

**Table 1.** Types and frequency of anomalies on the prosoma and opisthosoma in postembryos *Eratigena atrica*.

| Type of anomaly |  | Numer of individuals | % |
| --- | --- | --- | --- |
| oligomely | feeding appendages | 15 | 17.25 |
|  | walking appendages | 49 | 56.32 |
|  | feeding and walking appendages | 4 | 4.60 |
|  | spinnerets | 2 | 2.30 |
| schistomely | feeding appendages | 1 | 1.15 |
|  | walking appendages | 1 | 1.15 |
| symely | walking appendages | 1 | 1.15 |
| polymely | walking appendages | 1 | 1.15 |
| heterosymely | walking appendages | 1 | 1.15 |
| AP axis duplication |  | 6 | 6.89 |
| other abnormalities |  | 6 | 6.89 |
| Total |  | 87 | 100.00 |

Fig. 2A shows a spider with a duplication of the anterior prosoma, including the chelicerae. Two out of four chelicerae, located on the front of the prosoma on the left side (Ch1 and Ch2), had a regular shape and structure. On the right side of the prosoma, next to the Ch2, there were two supernumerary chelicerae (Ch3/4), whose basal segments were fused, while the cheliceral fangs were free. This individual had one mouth with a well-developed labium and lacked one walking leg on each side of the prosoma (oligomely).

Duplication of the anterior prosoma, including the chelicerae, can also be seen in the spider in Fig. 2B. This individual had four chelicerae on the front of the prosoma, three of which were fused. On the left side of the prosoma one chelicera was (Ch1) properly built, while on its right side was a large, massive appendage, composed of three chelicerae (Ch2/3/4), as evidenced by three movable cheliceral fangs. Only the basal segments of these chelicerae were fused along their entire length. The remaining prosomal appendages, i.e. a pair of pedipalps (P1 and P2) and four pairs of walking legs (L1-L4), had a regular structure. This individual had one mouth on the ventral side, with a well-developed labium between the gnathocoxae of the pedipalps P1 and P2.

A more profound change is presented in the individual in Fig. 2C. In this case the duplication of the anterior prosoma involved two body segments, i.e. the cheliceral and pedipalp ones, with obvious bicephaly. Both heads (HI and HII, on the left and on the right side of the prosoma) were complete, i.e. had fully developed chelicera with movable fangs (Ch1, Ch2, Ch3 and Ch4; HI and HII, respectively) and six-segmented pedipalps (P1, P2, P3 and P4; HI and H2, respectively). An extra appendage (Aa) between the gnathocoxa of the pedipalps P2 and P3, i.e. between the heads HI and H2 made this bicephalic case even more complicated. With only three articular surfaces, the appendage was significantly shorter and differed in structure from that of a properly constructed spider leg. It moved independently of the other prosomal legs. In addition, this individual had two mouths with well-marked labia and a set of walking legs (L1-L4).

Similar anomaly (duplication of the anterior prosoma involving two body segments with chelicerae and pedipalps) can also be distinguished in the bicephalic individual in Fig. 2D. One head (HI), located to the left of the body axis, was equipped with two chelicerae (Ch1 and Ch2) and two pedipalps (P1 and P2) and had one mouth with a well-developed labium. The other head (HII) also consisted of the chelicera and pedipalp metameres, but the chelicerae were fused at the basal segments (Ch3/4). The pedipalps (P3 and P4) were properly built, had a regular, six-segmented structure but the gnathocoxa of the pedipalp P4 was located more ventrally, directly behind the fused chelicerae Ch3/Ch4 in the place of the absent labium. This individual had four pairs of walking legs (L1-L4).

The spider in Fig. 2E had a more severe duplication of the prosoma. He had one fully developed head (HI) with two chelicerae (Ch1 and Ch2) and two pedipalps (P1 and P2) on the left side of the prosoma, while the other head (HII) was incomplete. It consisted only of the pedipalp segment with a pair of well-formed pedipalps (P3 and P4), whose gnathocoxae lay very close to each other, touching with their inner edges while the chelicerae were missing.

Immediately behind the pedipalps there was a very narrow, underdeveloped labium. In addition, between the two heads were four walking legs composed of 7 podomeres, which belonged to the first two duplicated segments of walking legs, (L1 and L2) On the ventral side of the prosoma, between the duplicated walking legs L2 a small, mobile protruberance (Pr) was visible. On the right and left side of the body behind the duplicated prosoma segments, there were four properly developed walking legs (L3-L4).

Duplication of the anterior prosoma was also diagnosed in the spider shown in Fig. 2F. The developmental defect in this individual consisted of the duplication of the chelicerae, pedipalps and the first two segments with walking legs. Additionally, heterosymely of the legs was observed. This individual had two complete heads (HI and HII), each equipped with a pair of chelicerae (Ch1, Ch2 and Ch3, CH4, respectively) and a pair of pedipalps (P1, P2 and P3, P4, respectively). Between pedipalps P2 (HI) and P3 (HII) there were four walking legs, which belonged to the duplicated segments L1 and L2. Two of them, i.e. L1, properly developed, flanked a heterosymelic complex of two walking legs (L2/2). The legs of the duplicated segment L2 were fused along almost their entire length up to the middle of the last podomere, i.e. the tarsus. Consequently, the heterosymelic complex had two short ends.

Additionally, this postembryo had only one well-developed labium between the gnathocoxae of the pedipalps P3 and P4 (HII), while the gnathocoxae of the pedipalps P1 and P2 (HI) were touching, and behind them was a protruberant oval structure, possibly corresponding to an abnormally developed labium. On the right and left sides of the body, behind the duplicated segments L1 and L2, there were two more pairs of walking legs of the segments L3 and L4. The AP axis duplication phenotypes from Figs. 2A-F are shown schematically in Figs 3A-F.

**Fig. 3.**
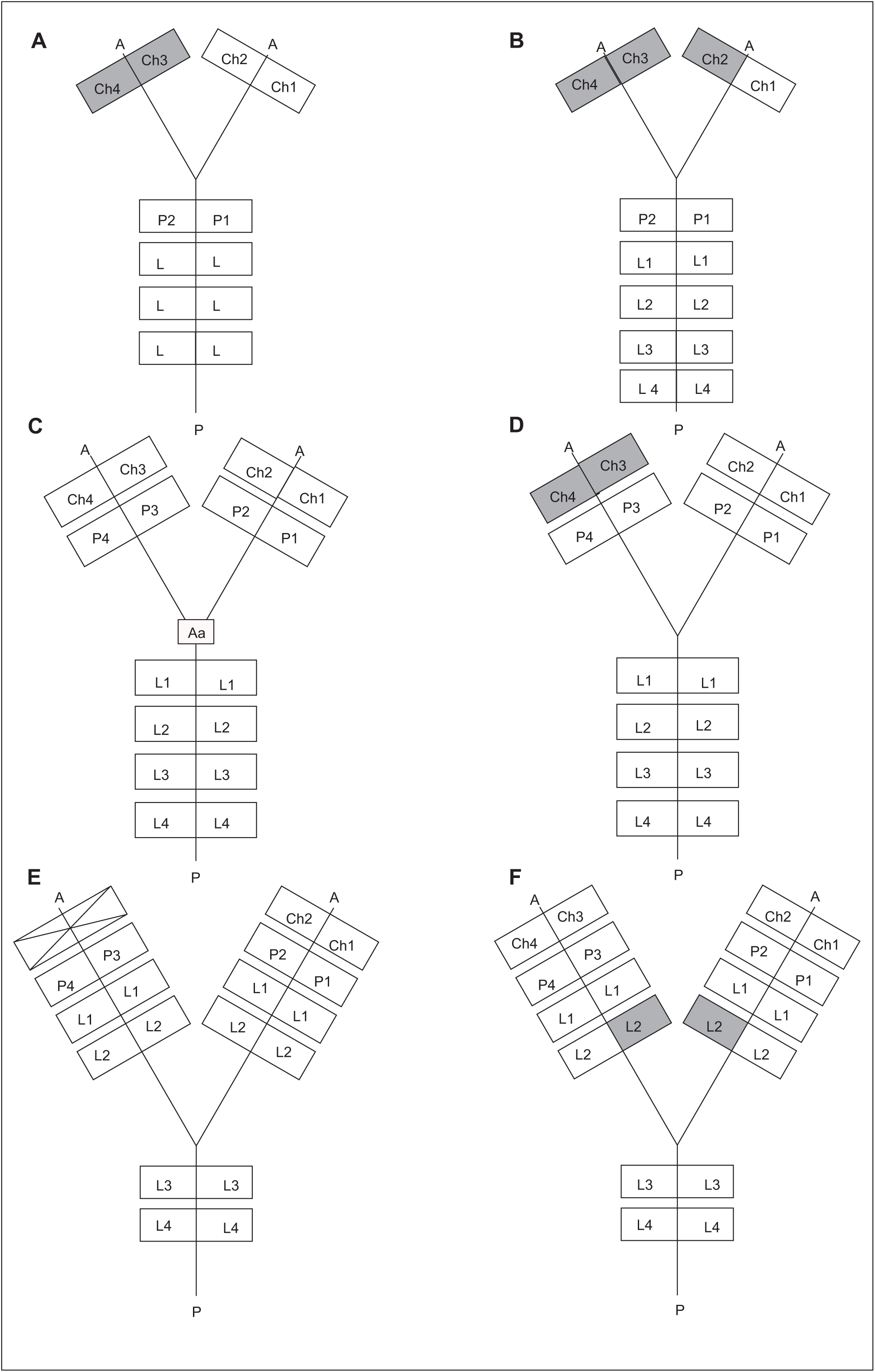
Six possible scenarios of deformities in postembryos shown in Figure 2A-F (ventral view). A:Duplication of the cheliceral segment with the fusion of two chelicerae (Ch3 and Ch4) and oligomely of walking legs in the postembryo in Fig. 2A; B: Duplication of the cheliceral segment with the fusion of three chelicerae (Ch 2, Ch3 and Ch4) in the postembryo in Fig. 2B; C: Duplication of the chelicerae and pedipalps with an additional hypophysis (Aa) between the pedipalps (P2 and P3) in the postembryo of Fig. 2C; D: Duplication of the cheliceral and pedipalpal segments with the fusion of two chelicerae (Ch3 and Ch4); E: Duplication of the prosoma up to the L2 segment accompanied by an absence of one pair of chelicerae in the postembryo in Fig. 2E; Duplication of the prosoma up to the L2 segment including simultaneous fusion of two walking legs (L2 and L2) in the postembryo in Fig. 2F; AP – anteroposterior body axis, dark squares indicate the fusion of appendages, a crossed out rectangle indicates a missing segment.

## DISCUSSION

No individuals with developmental defects were found in the control group, where embryo mortality was 8%. The results of teratological studies using variable temperatures in the first stages of embryonic development of *E. atrica* indicate that this factor not only significantly increases embryo mortality (38%), but also leads to morphological changes, which can be seen in postembryos leaving eggshells on their own. As in previous studies (e.g. Napiórkowska et al., 2016b, 2021), oligomely was the most common, affecting a significant percentage (80.46%) of teratogenically altered postembryos. The majority of the modified spiders lacked a walking leg or legs while only a few, feeding appendages or feeding appendages and walking legs. In two individuals oligomely affected the spinning apparatus. At present, owing to the availability of new tools and techniques, including the RNAi technique (e.g. Stollewerk et al., 2003; Schoppmeier and Damen, 2005; Oda et al., 2007; McGregor et al., 2008; Schwager et al., 2009 and Pechmann et al., 2011), we can safely assume that oligomely in spiders might be caused by the erroneous suppression or expression of segmentation patterning genes. Using this technique, Schwager et al. (2009) demonstrated that the reduction in the number of legs in the *Achaearanea tepidariorum* spider was related to the knockdown of the segmentation gene *hunchback* (*hb*) expression. The knockdown of its expression caused first instars hatched from *At-hb*^pRNAi^ egg sacs to have a reduced number of legs, i.e. two or three pairs instead of four. Another gene which was identified in the spider *Achaearanea tepidariorum* as a novel factor in anterior spider segmentation with a gap gene-like function is the *Distal-less* gene (*Dll*) (Pechmann et al., 2011). Pechman et al. demonstrated that the *Dll* gene of *A. tepidariorum* (*At-Dll*) is expressed unexpectedly early in the L1 region, before the formation of the germ band and the specification of the appendages. This early expression of *Dll* in the L1 region is necessary for the specification of the entire L1 segment, and is required for the subsequent activation of the segmentation genes in this region. The experiments of Pechmann et al. (2011) indicated that RNAi resulted in embryos lacking one entire leg-bearing segment (L1), and in a few embryos with permanently down-regulated At-Dll expression, the loss of two entire leg-bearing segments (L1 and L2). Other segmentation genes in spiders have also been studied. Oda et al. (2007) showed that parental RNAi against *At-Delta* resulted in defects not only in caudal lobe formation but also in the number of prosomal legs. The *Wnt8* gene also affects the development of body segments in spiders (McGregor et al., 2008). *At-Wnt8*^pRNAi^ embryos displayed a range of posterior phenotypes affecting leg-bearing segments 3 and 4 (L3 and L4). Based on the above results, we can assume that the reduction in the number of legs in *E. atrica* postembryos might have been caused by the erroneous suppression or expression of relevant segmentation genes. Teratological material also contained postembryos with other leg deformities, including schistomely, polymely, symely, heterosymely as well as significantly shortened or deformed legs, (classified among “Other abnormalities”, Table 1). There is strong possibility that at least some of these anomalies resulted from the loss of function of some developmental leg genes, also studied in the *Cupiennius salei* spider (Prpic et al., 2003; Prpic and Damen, 2004). For example, Prpic et al. (2003) proved that, among others, the genes *extradenticle* (*exd*), *homothorax* (*hth*), *dachshund* (*dac*) and *Distal-less* (*Dll*) have important roles during the formation of the proximal-distal (PD) axis of the *Cupiennius* leg. Therefore, we can conclude that all observed changes in the morphology of both the prosoma and its appendages may be the consequence of the stalling or erroneous expression of genes that model the body segmentation and determine the PD axis in appendages, following the application of alternating temperatures.

The exposure of embryos to alternating temperatures also led to and relatively rare anomalies, including pathological changes in the front of the prosoma. Six spiders affected in this way are presented in Fig. 2 A-F. Due to the nature of the deformities, they were classified as individuals with AP axis duplication (Table 1). The individuals in Fig. 2A and 2B had only the chelicera segment duplicated. Additionally, the basal segments of two or three chelicerae, respectively, were fused (Fig. 3A and 3B). It might seem that the presence of a supernumerary pair of chelicerae in a spider postembryo does not necessarily indicate bicephaly. However, previous histological studies (Napiórkowska et al., 2016c), where the CNS of such individuals was analyzed in detail, confirmed the presence of two brains in the prosoma. Therefore, we feel entitled to interpret this type of deformity as incomplete bicephaly. In two bicephalic individuals presented in Fig. 2C and 2D and Fig. 3C and 3D, the duplication of the cheliceral and pedipalpal segments was obvious. Additional changes included the presence of a shortened appendage between the two heads (Aa) and the fusion of the basal cheliceral segments (Ch3/4) on one of the heads. Based solely on its morphology, it is difficult to determine whether the appendage (Aa) was a shortened pedipalp or a modified supernumerary walking leg, and whether this change was related to the duplication of the AP body axis. It can be assumed that thermal shocks disturbed the expression of segmentation genes as well as the genes responsible for leg structure, which resulted in the formation of half of the metamer along with a modified appendage. At the same time the fusion of the basal cheliceral segments on HI in the individual shown in Fig. 2D, 2A and 2B remains unexplained, with no information in the literature about the mechanisms of appendage fusions in spiders. We cannot dismiss the possibility that this occurs as early as when the appendage buds are formed and the process is related to abnormal cell movements or the elimination of some of them. In the study two individuals had a more profound duplication of the AP axis (Fig. 2E and 2F; Fig. 3E and 3F): apart from the head segments, the segments of the L1 and L2 walking legs were also duplicated. In the spider in Fig. 2E, the defect affected only the pedipalp segment while the cheliceral segment was not duplicated, and in the spider in Fig. 2F, an additional morphological complication, i.e. the fusion of the walking legs L2/L2)(heterosymely) was observed.

At present, the mechanism of bicephaly cannot be determined owing to the lack of systematic research. We can only assume that this malformation is cellular and might be related to the displacement or dislocation of the cumulus, visible as a little bulge on the egg surface in early spider embryogenesis (Chaw et al., 2007; Wolff and Hilbrant, 2011; Mittmann and Wolff, 2012). It is a group of mesenchymal cells that function as a migratory cell cluster to break the radial symmetry of the germ-disc. Akiyama-Oda and Oda (2003) found that the cumulus of *Achaearanea tepidariorum* (now *Parasteatoda tepidariorum*) is a site of decapentaplegic (dpp) signaling. The epithelial cells that overlie the path of cumulus migration extend cytonemes to the cumulus and respond to the dpp signaling. In response to cumulus migration and dpp signaling, the embryonic germ disc breaks radial symmetry. The direction of cumulus movement allows one to predict the orientation of the future AP and DV axes (Oda et al., 2020). The functional importance of the cumulus in early patterning of the spider embryo was demonstrated by Holm (1952), who used embryos of the spider, *Agelena labyrinthica*. He extirpated the cumulus during migration in *Agelena* embryos. This manipulation resulted in embryos showing persistent radial symmetry with no typical germ band formed. Holm transplanted too a part of the cumulus to the opposite side of the same embryo and showed a huge variety of axis duplication phenotypes that can occur if cumulus material is placed at different regions within the germ-disc. His manipulation resulted in doubling of bilaterally symmetric body patterns in a single egg, producing two separate sets of AP and DV axes. This experiment provided strong evidence of the cumulus acting as an organizer capable of inducing an additional set of axes defining bilateral body symmetry.

Experiments by Holm (1952) inspired other researchers to obtain twinned embryos by transplantation of the cumulus to ectopic sites. Oda et al. (2020) obtained twins by transplanting cumuli between sibling embryos in jumping spider, *Hasarius adansoni*. These manipulations showed that a grafted cumulus appeared to induce ectopic extraembryonic tissue, and two sets of body axes were formed between the intact and ectopic extraembryonic regions. Thus, according to the authors, the cumulus seems to function as an organizer of bilateral symmetry defined by two body axes and may be a common feature of the spiders. A similar study was conducted by Pechmann (2020). He performed cumulus grafting in *Acanthoscuria geniculata* embryos at mid to late stage 5. The ectopic cumulus was always placed within the germ-disc opposite to the endogenous cumulus, which was already migrating towards the rim of the disc. The cumulus material from the donor embryo was clamped between the germ-disc and the vitelline membrane of the acceptor embryo. The cumulus of the donor embryo regularly fused with the ectoderm of the germ disc of the acceptor embryo. As cumulus material from the donor embryo sometimes detached from the germ-disc of the acceptor embryo, not all embryos showed an axis duplication phenotype. In the end, he obtained 12 embryos that showed a complete or partial axis duplication phenotype, which means that twinning was more or less perfect. According to Pechmann (2020), the fusion of prosomal segments and imperfect twinning of embryos was probably due to the fact that grafted cumulus material was not big enough or that the fusion of the grafted material did take place at the wrong place and time. In these cases the developing embryos shared a single wild type looking opisthosoma. Thus, based on the results of Oda et al. (2020) and Pechmann (2020), we believe that the observed imperfect twinning of postembryos resulted from disturbances in the formation of cumulus material or its displacement or dislocation. The importance of cumulus in modeling the body axis not only in spiders, but also in myriapods is confirmed by the results of Janssen (2013). He observed abnormally developing embryos of the common pill millipede *Glomeris marginata*, all of which represent cases of *Duplicitas posterior*, which means that the posterior body pole was duplicated. He recorded four classes of abnormalities: (1) symmetrically duplicated germ bands that share a single anterior pole, (2) asymmetrically duplicated germ bands that share a single anterior pole; (3) duplicated posterior germ bands without anterior pole; (4) duplicated posterior germ bands with a rudimentary anterior pole. According to Janssen (2013), a plausible reason for the occurrence of *D. posterior*-type embryos could be the formation of two posterior organization centers (cumuli) instead of only one, as the result of early development at excessive temperatures. In our study, it was alternating temperatures that led to the phenotypes of front body duplication (*Duplicitas anterior*). This assumption may be supported by the fact that spiders are ectotherms and temperature has a significant impact on their lives, both in the physiological and ecological context. Temperature not only affects the growth and survival rate of juvenile stages, but also determines the course and duration of embryogenesis (Li, 1995 2002; Li and Jackson, 1996; Hanna and Cobb, 2006; Napiórkowska et al., 2018). Since *E. atrica* embryos were exposed to temperature fluctuations that may impact orphogenetic processes, we assume that all recorded body anomalies and relatively high embryo mortality were the consequence of using the variable temperature protocol, and the DA anomaly was not caused by genetic mutations (could not be inherited) but resulted from disturbances in early embryonic development (in the formation and migration of the cumulus).

## MATERIALS AND METHODS

In our teratological experiment we used embryos of the synanthropic spider *Eratigena atrica* (C.L. Koch, 1843) of the Agelenidae family. In September, 28 sexually mature females and 16 males were collected in the vicinity of Toruń and Chełmża (Poland) to establish a laboratory culture. Spiders were kept in a dark room with the temperature of 21°C and a relative humidity (RH) of 70%. Each spider was placed in a 250 cm3 ventilated glass container and fed twice a week with Tenebrio molitor larvae. In the first days of October, a male ready for mating was introduced to each female. This procedure was repeated one week apart to ensure that all females were inseminated. Since the females outnumbered males, each male mated with multiple females, thus becoming the father of several mothers’ offspring. First egg sacs were laid after a few weeks, followed periodically by additional egg sacs, averaging 8 egg sacs per female. They were removed from the culture vessels, cut open to obtain eggs, which were then randomly dipped in paraffin oil (three eggs from each egg sac) to confirm their fertilization. Counted embryos were divided into two groups, control and experimental.

Embryos from the control group were incubated at 22°C and 70% RH until postembryo hatching. Embryos from the experimental group were exposed to temperatures of 14°C and 32°C (70% RH), changed every 12 hours. Alternating temperatures were applied for 10 days, until segments of the prosoma appeared on the germ band and limb buds appeared on these segments [comparable to Stage 9 in the trechaleid *Cupiennius salei* (Keyserling, 1877) (Wolff and Hilbrant, 2011); Stage 8.2 in the theridiid *Parasteatoda* (formerly *Achaearanea*) *tepidariorum* (C.L. Koch, 1841) (Mittmann and Wolff, 2012) as determined by dipping randomly selected embryos in paraffin oil. Further incubation of the experimental embryos was carried out in the same conditions as the control ones. After hatching, all postembryos were examined for the presence of anomalies on the prosoma and the opisthosoma.

Individuals with the most interesting deformities, including bicephaly, were photographed using a light microscope (Axcio Lab A1). Images were recorded using digital camera (Axiocam 105 color, Carl Zeiss) and a computer system running Zen software (Version 2.3, blu edition).

## Data availability

All relevant data can be found within the article.

